# Non-fatiguing physical exercise induces sustained changes in motor unit discharge patterns and estimated synaptic inputs to motoneurons

**DOI:** 10.64898/2026.11.12.765939

**Authors:** Titouan Masson, Valentin Goreau, François Hug, Thomas Cattagni

## Abstract

Non-fatiguing exercise modulates neural excitability in motor pathways controlling both exercised and unexercised muscles. Whether these effects also alter the synaptic control of spinal motoneurons remains unclear. This study investigated how acute non-fatiguing exercise influences motor unit discharge patterns and the estimated contributions of neuromodulatory input and excitation-inhibition balance, depending on muscle activation during exercise.

Fourteen participants completed two 30-minute rowing conditions involving either the upper body or the full body. High-density surface electromyography was used to assess tibialis anterior motor unit discharge properties before and up to one hour after exercise. The contributions of neuromodulatory input and excitation-inhibition balance to motor unit discharge were estimated using paired motor unit analysis and geometric characteristics of discharge patterns.

At matched torque, peak motor unit discharge rates increased following both exercise conditions, with effects persisting for up to 45 minutes. However, the estimated contributions of synaptic inputs differed between conditions. Following upper-body exercise, when the tibialis anterior was minimally activated, changes in motor unit discharge patterns suggested an enhanced contribution of neuromodulatory input, with no clear changes in excitation-inhibition balance. In contrast, the analyses following full-body exercise suggested a lower contribution of inhibition with respect to excitation, without clear evidence of enhanced neuromodulatory input. These findings suggest that similar increases in motor unit discharge rates following non-fatiguing exercise can be associated with different contributions of synaptic inputs.

Overall, our data suggest that acute non-fatiguing exercise induces sustained changes in motor unit discharge patterns, even in muscles minimally involved in the task. The estimated contributions of neuromodulatory input and excitation-inhibition balance to these changes appear to depend on muscle activation during exercise, providing new insights into the neural regulation of spinal motoneuron activity following non-fatiguing exercise.

## INTRODUCTION

Acute physical exercise can rapidly and transiently modulate motor system excitability. At the cortical level, non-fatiguing aerobic exercise increases intracortical excitability and may also modulate corticospinal excitability (Singh *et al*., 2014; Yamazaki *et al*., 2019; Youssef *et al*., 2024; Harroum *et al*., 2025). Importantly, some of these effects are also observed in neural pathway controlling muscles not directly involved in the exercise, suggesting a diffuse central modulation alongside muscle-level adjustments (Singh *et al*., 2014. Yamazaki *et al*., 2019; Youssef *et al*., 2024). Exercise can also modify the Hoffmann reflex, suggesting changes within spinal sensorimotor pathways (Motl et *al.,* 2003; Motl *&* Dishman, 2003). However, because the Hoffmann reflex depends on several mechanisms, these changes cannot be specifically attributed to motoneuron excitability. Thus, whether the diffuse central effects of acute exercise extend to spinal motoneurons, and how they interact with local muscle activation, remains unclear.

Described as the final common pathway of the motor system (Sherrington, 1906), spinal motoneurons translate these central adjustments by integrating both ionotropic and neuromodulatory synaptic inputs. Ionotropic excitatory and inhibitory inputs shape membrane potential and directly influence motoneuron recruitment and discharge behavior (Powers *&* Binder, 2001; Pierrot-Deseilligny *&* Burke, 2012). Neuromodulatory input, particularly serotonergic projections, modulate motoneuron intrinsic properties through G-protein-coupled receptors that facilitate persistent sodium and L-type calcium currents, generating persistent inward currents (PICs) (Heckman *et al*., 2008; Powers *&* Heckman, 2015). PICs amplify and prolong excitatory synaptic inputs, enabling motoneurons to sustain firing rate despite reductions in net ionotropic excitatory drive (Gorassini *et al*., 2002; Lee *&* Heckman, 1999). They play a major role in determining motoneuron input-output gain and force production (Heckman *et al*., 2005).

Acute exercise is known to modulate the monoaminergic neuromodulatory system. In animal models, 15-30 minutes of physical exercise increases the activity of serotonergic neurons in the dorsal raphe nucleus (Otsuka *et al*., 2016; Morikawa *et al*., 2021), while 20-60 minutes of physical exercise increases brain serotonin release, with effects persisting for 20-30 minutes after exercise cessation (Meeusen *et al*., 1996; Gómez-Merino *et al*., 2001). In humans, indirect evidence suggests that exercise may also enhance neuromodulatory influences on spinal motoneurons. Indeed, upper-limb contraction induces an increased estimates of PICs contribution to lower-limb motoneuron firing (Orssatto *et al*., 2022; Ugliara *et al*., 2026). Furthermore, neuromodulatory input proxy was increased after hand grip task (Ugliara *et al*., 2026) and during arm-cycling task (Nazaroff *et al*., 2026). Conversely, repeated maximal muscle contractions reduce PICs estimates in the exercised muscle (Mackay *et al*., 2023), potentially due to fatigue-related local inhibitory feedback (Amann *et al*., 2015) or excessive serotonin release (Mackay *et al*., 2023; Cotel *et al*., 2013). Together, these findings suggest that exercise-induced changes in motoneuron excitability may reflect both systemic neuromodulatory effects and mechanisms at muscle level. However, how these influences interact following non-fatiguing exercise, particularly depending on whether a muscle is directly involved or not, remains unknown. In addition, although animal studies suggest that exercise-induced neuromodulatory effects persist after exercise cessation, their time course and dependence on muscle involvement remain poorly understood in humans.

This study aims to determine the time course of changes in motor unit discharge properties over the first hour following acute, non-fatiguing exercise and to investigate whether the mechanisms underlying these changes differ according to muscle involvement. To address this, we assessed tibialis anterior motor unit discharge properties and estimated the contributions of neuromodulatory input and excitation-inhibition balance at multiple time points before and after exercise, comparing conditions where the muscle was moderately involved (i.e., full-body exercise) or minimally involved (i.e., upper-body exercise). We hypothesized that motoneuron peak discharge rates would increase after exercise, with effects persisting for several minutes regardless of exercise condition. Specifically, we expected this facilitation to be driven primarily by systemic neuromodulatory input when the muscle was minimally involved. Conversely, when the muscle was moderately involved, we hypothesized that the increased discharge rate would instead be sustained through local adaptations in excitation-inhibition balance.

## METHODS

### Participants and ethical approval

Fourteen participants (21.6 ± 6 years, 1.81 ± 0.1 m, 72.1 ± 13.7 kg, 2 women) were recruited. Participants had to report no lower limb injuries in the previous 6 months. They were instructed not to train or engage in physical activity for 24 hours before the evaluation. The study protocol was approved by the local ethics committee (CERNI, Nantes Université, IRB: IORG0011023). All procedures conducted in this study adhered to the requirements of the Declaration of Helsinki (last revised in 2024). The experimental protocol and any potential discomfort were explained to the participants and written informed consent was obtained prior to testing.

### Experimental setup

Evaluations were performed on the tibialis anterior of the participant’s dominant leg (the one used to kick a ball) during isometric dorsiflexor contraction. Participants were tested in a seated position with the hip angle at 80° of hip flexion (neutral position = 0°), the knee angle at 0° (neutral position), and the ankle angle at 10° of plantar flexion (neutral ankle position = 0°). Participant’s foot was securely attached to the footplate. The lateral malleolus was aligned with the ergometer’s axis of rotation. Participants were instructed to perform isometric voluntary contractions consisting of a 10-second ramp-up phase and a 10-second ramp-down phase, peaking at 20% of MVC. To perform these triangular ramp task, participants were following real-time torque feedback displayed on a computer screen. These contractions were used to perform quasi-geometric approach (Beauchamp *et al*., 2023) and to calculate ΔF using the paired motor unit method (Gorassini *et al*., 2002).

#### Mechanical recordings

Torque measurement during isometric dorsiflexor contraction were assessed using a custom-made pedal equipped with strain gauges (Société Doerler–Vandoeuvre, France) and an amplifier (PM Instrumentation, model 1300B, amplification: 8 V = 200 Nm) developed by an engineering school (University Institute of Technology Mechanical Engineering, Dijon, France) (Fig. 1A). The strain gauges output signal was collected at 2 048 Hz using the multi-channel HDsEMG acquisition system (EMG-Quattrocento, 400 channel EMG amplifier; OT Bioelettronica).

#### Electromyography recordings

HDsEMG signals were recorded from tibialis anterior muscle during isometric dorsiflexor contraction. 4 two-dimensional adhesive grids of 64 electrodes were placed [13 x 5 gold-coated electrodes with one electrode missing in one corner, interelectrode distance: 4 mm (GR04MM1305, OT Bioelettronica, Italy)] (Fig. 1A). Before applying the electrode grids, the skin was shaved and cleaned with an abrasive pad. The adhesive grids were held on the skin with semi-disposable bioadhesive foam layers (SpesMedica, Battipaglia, Italy). Skin-electrode contact was established by filling the cavities of the adhesive layers with conductive paste (SpesMedica, Battipaglia, Italy). A reference electrode (Kendall Medi-Trace, Canada) was placed over the patella and a ground electrode (strap dampened with water) was placed around the ankle of the tested leg. EMG signals were recorded in monopolar mode, band-pass filtered (10-500 Hz), and digitized at a sampling rate of 2 048 Hz using a multi-channel HDsEMG acquisition system (EMG-Quattrocento, 400 channel EMG amplifier; OT Bioelettronica). The data were acquired using OT BioLab+ software (OT Bioelettronica, Torino, Italy).

### Experimental protocol

This study included three randomized experimental sessions performed on different days, performed at the same time of the day within each individual. A session was dedicated to the assessment of neuromuscular fatigue in tibialis anterior after upper-body and full-body rowing. Two other sessions were designed to assess motor unit behaviour during torque-matched isometric dorsiflexion tasks performed before and throughout the hour following either upper-body rowing or full-body rowing. This design allowed us to compare the time course of motor unit behaviour after both exercise conditions (i.e., moderately or minimally involved muscle).

#### Fatigue assesment session

Preliminary tests had been performed to estimate the average activation level of the tibialis anterior muscle during both conditions. Mean activation during the upper-body rowing was 1.7 ± 1.1% of maximal voluntary contraction (MVC), whereas the full-body rowing elicited a mean activation of 14.7 ± 6.3 % of MVC. To ensure that neither condition induce fatigue, the following fatigue assessment session was designed. First, participants began with a standardized warm-up, consisting of two 5-second dorsiflexor contractions at intensities of 50%, 70%, and 90% of their self-estimated MVC, with a 20-second rest period between each contraction. After a 2-minute rest, participants performed two 5-second dorsiflexor MVCs on the custom-made pedal, with a 2-minute rest period between contractions. If the difference between the two contractions exceeded 5%, an additional MVC was performed. During each MVC attempt, participants crossed their arms over their chest and received real-time visual feedback of the torque signal, as well as strong verbal encouragements. Second, participants performed 30 minutes of physical exercise on an air rowing machine (RowErg Standard Concept2, Morrisville, USA), involving either full-body rowing or only upper-body rowing. In the latter condition, participants rowed with feet anchored to the ground and knees maintained at approximately 90° of flexion. During both rowing conditions, participants were instructed to maintain a target heart rate corresponding to 65% of their age-predicted maximal heart rate, calculated as: (220 - age) x 0.65 beats per minute (bpm) (Astrand *&* Ryhming, 1954). To provide continuous visual feedback of heart rate, participants wore a pectoral heart rate monitor (Polar H10, Polar Electro Oy, Finland). The mean power output in Watt (W) was recorded from the air rowing machine at the end of each exercise session. Third, participants switched to the custom-made pedal to perform two MVCs under the same conditions as step one, allowing exactly one minute of transition after the rowing exercise. The entire procedure (rowing bout and subsequent MVC measurements) was then repeated for the remaining condition.

**Figure 1.**
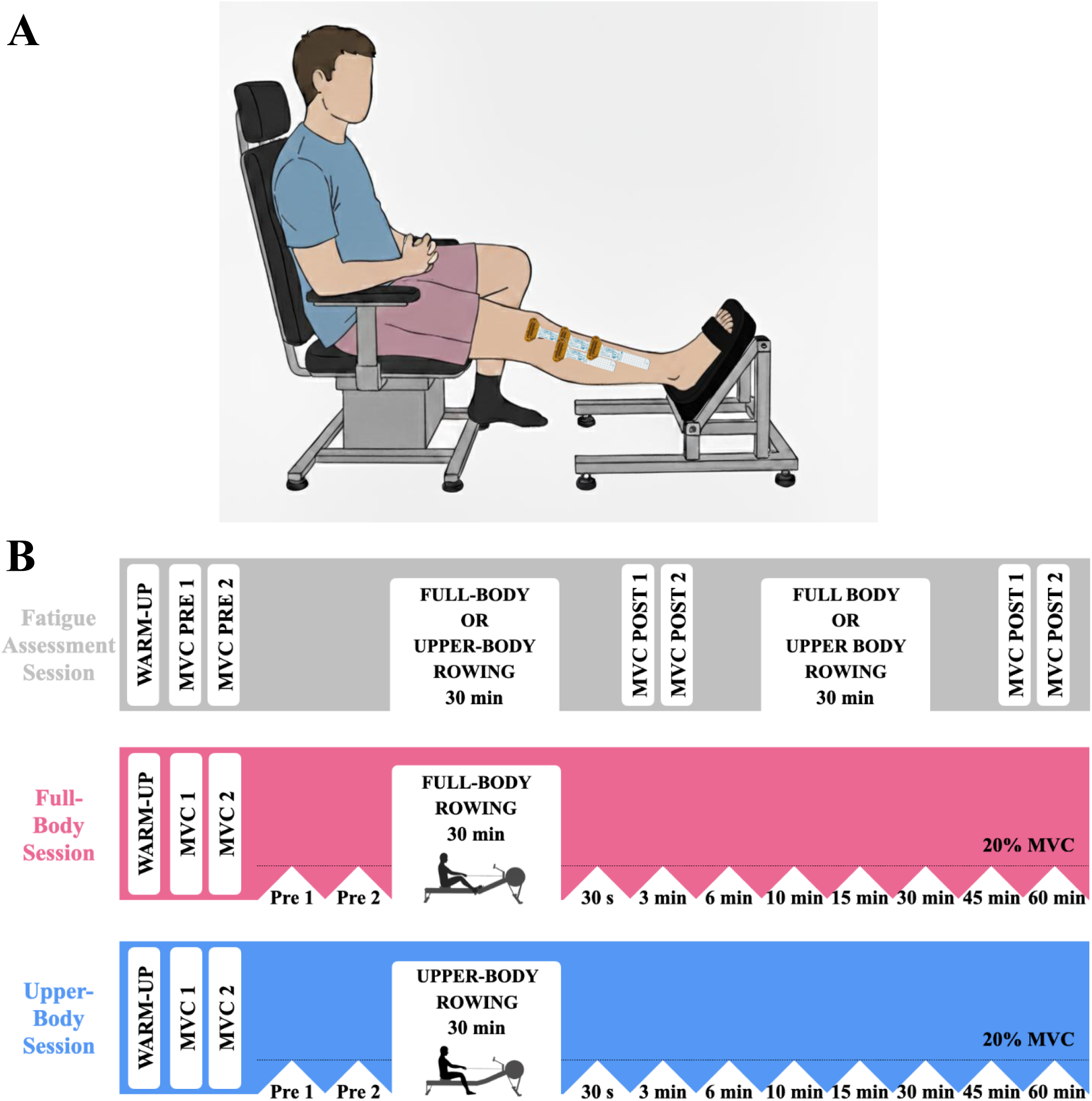
Experimental protocol. Participants were tested in a seated position with the hip angle at 80° of hip flexion (neutral position = 0°), the knee angle at 0° (neutral position), and the ankle angle at 10° of plantar flexion (neutral ankle position = 0°) (*A*). Participant’s foot was securely attached to the footplate. The lateral malleolus was aligned with the ergometer’s axis of rotation. Four high density surface electromyography grids (256 electrodes in total) were positioned over the tibialis anterior muscle. Participants were tested during three different sessions (*B*). During fatigue assessment session (in grey), after a warm-up, participants performed both rowing condition in a random order and realized maximal voluntary contractions (MVCs) before and after each session to estimate fatigue generation. During full-body (in pink) and upper-body (in blue) sessions, participants performed a warm-up, followed by two MVCs. Then, they performed isometric submaximal dorsiflexor contractions before and for 60 minutes after an exercise session involving either full body or the upper body only. Each contraction represents a triangular isometric contraction lasting 20 seconds (10 seconds ramp-up and 10 seconds ramp-down), peak level corresponding to 20% of MVC.

#### Neuromuscular assessment sessions

The other two sessions were used to collect HDsEMG data as follows. First, participants began with a standardized warm-up (as described in fatigue assessment session). Second, participants were familiarized with isometric dorsiflexor contraction during triangular ramp task. Once they were able to accurately follow the torque feedback, participants performed two triangular contractions with a 3-minute rest interval between each trial. Trials were excluded and repeated if the torque trajectory was not closely matched, for example, a sharp increase or decrease in the torque signal. Third, participants performed 30 minutes of physical exercise on an air rowing machine involving either full-body or upper-body mobilization (as described in the fatigue assessment session). This 30-minute duration was selected based on established human paradigms demonstrating that 20 to 30 minutes of aerobic exercise is sufficient to induce systemic neuromodulation, as evidenced by modulated intracortical networks (Singh et al., 2014; Yamazaki et al., 2019). Fourth, to characterize the time-course of this systemic neuromodulatory input and test our hypothesis, participants performed isometric voluntary triangular contractions at the following post-exercise time points: 30 seconds, 3 minutes, 6 minutes, 10 minutes, 15 minutes, 30 minutes, 45 minutes, and 60 minutes (Fig. 1B).

### Data analysis

#### HDsEMG Decomposition

We used the convolutive blind source separation method (Negro et al., 2016) implemented in an open-source software (Muedit; Avrillon *et al*., 2024) to decompose the EMG signals into motor unit spiking activity. The automatic decomposition was performed on the monopolar EMG signals (noisy channels excluded) from one contraction per timing in each condition. Then, we manually edited all the motor unit spike trains from each triangular contraction following previously published procedures (Del Vecchio *et al*., 2020; Hug *et al*., 2021). The editing process involved an iterative approach, discarding detected peaks that result in erroneous firing rates (outliers) and adding missed firing times that are clearly distinguishable from the noise. The motor unit pulse trains were recalculated with updated motor unit filters.

#### Tracking of the motor units across contractions

To ensure the comparison of the same motoneurons between contractions within each condition, it would have been relevant to match motor units. However, when matching motor units across time in each condition, the available motor unit pool was reduced, with a loss of 72% and 77% of the initially identified motor units in the upper-body and full-body conditions, respectively. As this dramatically reduced the dataset, we reported the unmatched motor unit in the present paper. Data from matched motor units are reported as supplemental material. Importantly, the results obtained from the matched motor unit analysis were consistent with those obtained from the unmatched motor unit analysis.

#### Motor unit discharge analysis

For each motor unit, the instantaneous discharge rates were smoothed using support vector regression to create continuous estimates (Beauchamp *et al*., 2022). The maximum value from the smoothed discharge rate of each motor unit was considered as the peak discharge rate. The recruitment threshold of each motor unit was defined as the torque at its first discharge.

We used a quasi-geometric approach to assess how much the motor unit discharge rate deviated from a linear relationship with torque output during the ramp-up phase of the contractions (Fig. 2). In this approach, the smoothed discharge rate of each motor unit was plotted against the torque generated during a contraction. The largest perpendicular distance between this discharge rate and a hypothetical linear increase from recruitment to peak discharge rate was quantified, and this distance was defined as brace height. To ensure that brace height reflects only the relative deviation from a linear pattern and to eliminate any bias from changes in peak discharge rates, brace height was normalized. Specifically, it was normalized to the height of a right triangle, where the hypotenuse runs from the point of recruitment to the peak discharge rate. Previous studies, using simulations of motoneuron behavior, have shown that brace height is highly sensitive to changes in neuromodulatory input (i.e., PICs activation) and negligibly sensitive to changes in excitation-inhibition balance (Beauchamp *et al*., 2023; Chardon *et al*., 2024).

In addition to brace height, we calculated the slopes of both the initial acceleration phase and the subsequent attenuation phase of the motor units discharge rate (Beauchamp *et al*., 2023). Acceleration is characterized by a sharp increase in motor unit discharge rate right after recruitment, which is likely due to the activation of PICs (Revill *&* Fuglevand, 2017). This phase is followed by an attenuation phase, where the increase in discharge rate slows down as PICs conductance continues to rise and potassium currents are activated (Li *&* Bennett, 2007). The rapid acceleration in motor unit discharge rate during acceleration is thought to be driven by PICs, making the acceleration slope an indicator of PICs amplification, however, it has been shown in computational models that this indicator is influenced by inhibition. The attenuation phase, on the other hand, is associated with the excitation-inhibition balance (i.e., reciprocal pattern or proportional pattern) (Beauchamp *et al*., 2023; Chardon *et al*., 2024), and may thus, in combination with brace height and ΔF metrics, allow to get further insights into the contribution of neuromodulatory input and excitation-inhibition balance to motor units discharge patterns.

Motor units that showed a negative slope during the acceleration phase or had a negative brace height prior to normalization were excluded from further analysis. Quasi-geometric approach could unintentionally create artificial acceleration and attenuation phases if the true peak discharge rate had not been reached during the contraction. To avoid this, motor unit discharge patterns were excluded if the duration of discharge before peak discharge rate was shorter than 3.5 seconds. This threshold was chosen because it enables a full acceleration phase and a sufficient attenuation phase to perform slope analysis.

**Figure 2.**
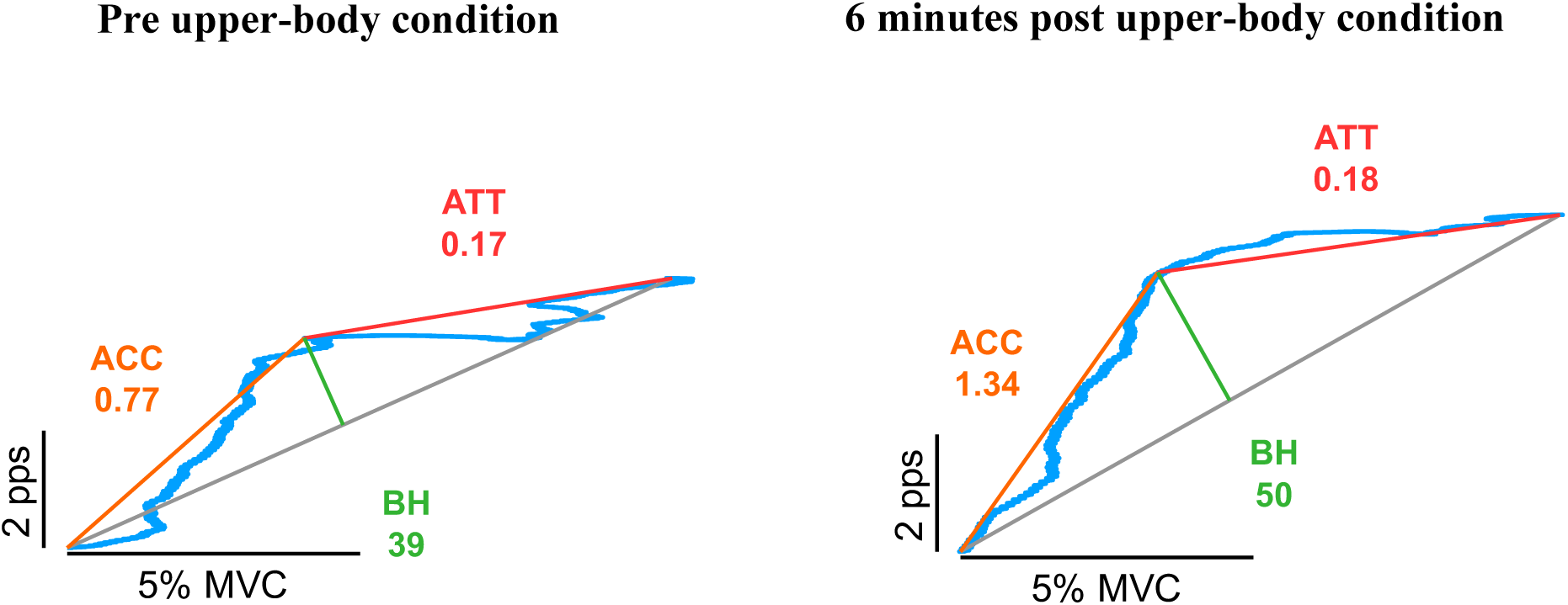
Example of motor unit discharge rate analysis using the geometric approach. This approach calculates brace height (BH, %), acceleration (ACC, pps/%MVC), and attenuation (ATT, pps/%MVC), which were used respectively to estimate the contribution of the neuromodulatory drive to motor unit discharge, estimate of persistent inward currents amplification on the discharge, and the pattern of inhibition. The same motor unit discharge rate and indicators are shown before (left chart) and 6 minutes after upper-body condition exercise (right chart).

Then, the prolongation effect of PICs was estimated using the paired motor unit method (Gorassini *et al*., 2002). Motor units with a lower recruitment threshold (control units) were paired with motor units with a higher recruitment threshold (test units). ΔF was calculated as the change in discharge rates of the control motor unit from the moment of recruitment to the moment of derecruitment of the test unit (Gorassini *et al*., 2002) (Fig. 3). In line with previous work (Hassan *et al*., 2020), the criteria for including a pair of motor units were as follows: 1) the test motor unit should discharge for at least 2 seconds, 2) the test motor unit should be recruited at least 1 second after the control motor unit to ensure full activation of PICs, 3) the test motor unit should be derecruited at least 1.5 second before the control motor unit to prevent overestimation of ΔF, 4) the control unit modulated its discharge rate by ≥ 0.5 pps while the test unit was active (Stephenson & Maluf, 2011), 5) a coefficient of determination (r^2^) > 0.7 should be observed between the smoothed discharge rate of the test and control motor unit (Hassan *et al*., 2020). ΔF values are presented as “unit-wise” averages for all suitable test-control unit pairs, thus reducing the number of ΔF values to one per test unit.

**Figure 3.**
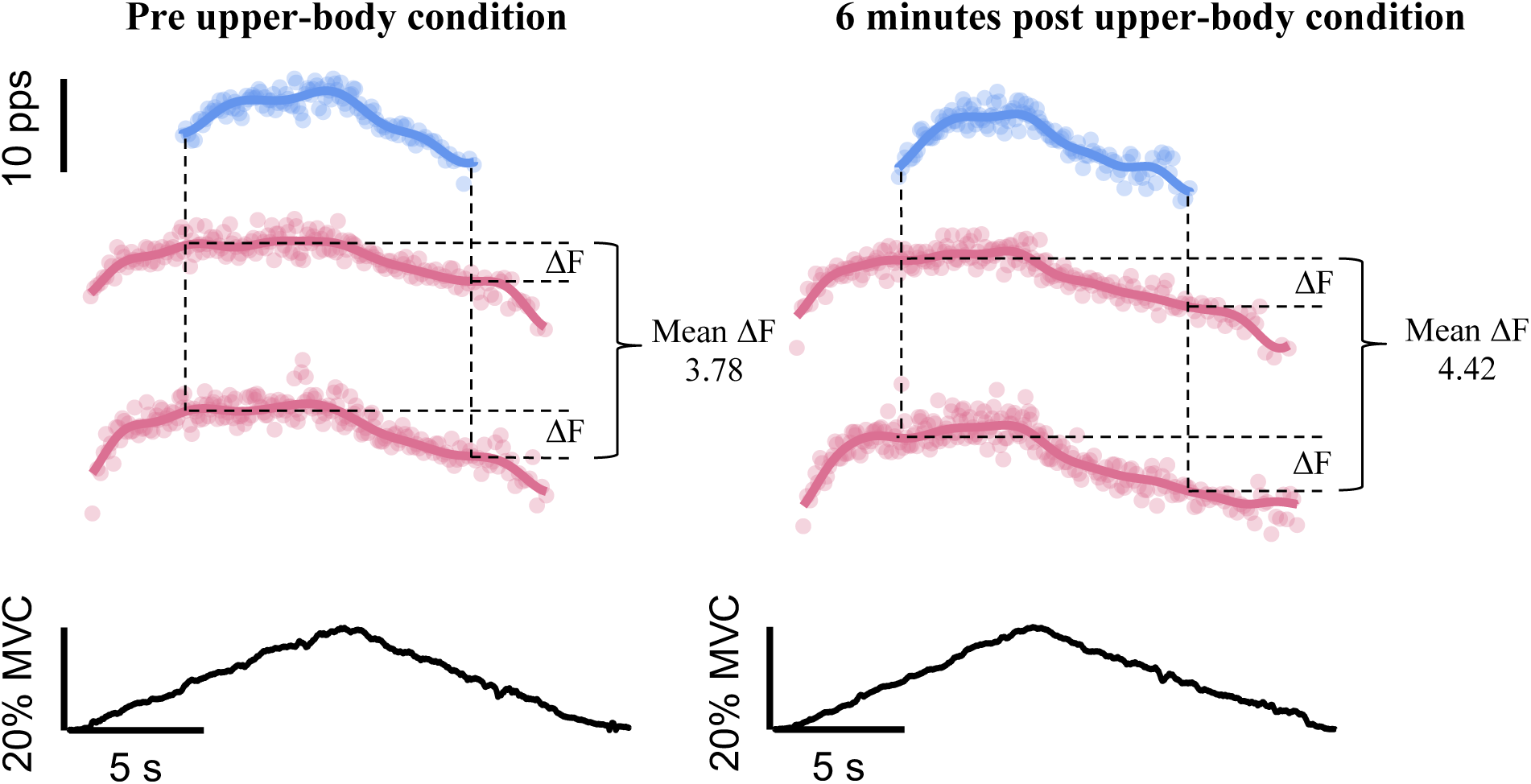
Example of estimate of persistent inward current prolongation analysis using the paired motor unit method (ΔF, pps) In this example, the same test motor unit’s smoothed discharge rate is shown in blue, while the control unit’s smoothed discharge rates are shown in pink. Since several motor units can be used to estimate ΔF, the average ΔF value across all units was used for the analysis. This figure represents real data from our study. Paired motor unit analysis on a test motor unit is presented with data from before upper-body condition (left chart) and from 6 minutes after upper-body condition (right chart).

### Statistical analysis

To assess changes in motor units discharge characteristics across timings for each condition, we applied a linear mixed-effects model to each motor units discharge metric (i.e., recruitment threshold, peak discharge rate, brace height, acceleration, attenuation and ΔF), incorporating data from individual motor units. The model included conditions (full-body and upper-body) and timings (pre, 30 seconds, 3 minutes, 6 minutes, 10 minutes, 15 minutes, 30 minutes, 45 minutes and 60 minutes) as fixed effects. For random effects, we added a random intercept and a random slope for each participant to account for the variability in the timing effect between participants [e.g., ΔF ∼ Condition*Timing + (Timing | Participant)]. In cases where the model failed to converge, we simplified the random structure by removing the random slope for each timing. This adjustment preserved the key random structure that accounts for the effect of timing across participants. Linear mixed-effects model analysis was performed using R (v.4.4.1, R Core Team 2021, R Foundation for Statistical Computing, Vienna, Austria), with the lmerTest package (Kuznetsova *et al*., 2017) to fit the models. For each model, we graphically checked that the assumptions of homoscedasticity, normality and independence of residuals were met through graphical evaluations. When significant effects were detected, we applied Dunnett’s post-hoc test to compare each post-exercise timing with the pre-exercise timing, used as the reference. Additionally, we calculated estimated marginal mean differences and standardized differences, along with 95% confidence intervals, using the emmeans package. Significant difference was accepted at p < 0.05.

## RESULTS

### Physical activity parameters

Participants mean age was 21.6 ± 6.0 years, therefore the average target heart rate for the rowing exercises in both sessions was 129.0 bpm. The mean heart rate was 123.1 ± 10.0 bpm during upper-body condition and 128.8 ± 4.9 bpm during full-body condition. During the fatigue assessment session, the mean heart rate was similar to values from experimental sessions: 123.1 ± 8.7 bpm in upper-body condition and 128.7 ± 4.9 bpm in full-body condition. The mean power output was 49.2 ± 17.4 W during upper-body condition and 97.4 ± 27.4 W during full-body condition. Pre-exercise mean MVC was 37.4 ± 7.3 N.m. After full-body condition, the mean MVC was 37.5 ± 5.6 N.m, which represent 100.3% of the pre-exercise MVC. After upper-body condition, the mean MVC was 37.1 ± 5.9 N.m, representing 99.2% of the pre-exercise MVC. These results highlight that these exercises did not generate measurable neuromuscular fatigue.

### Motor unit decomposition

In upper-body condition, 31.6 ± 10.7 motor units per participant were identified in pre-exercise and an average of 21.7 ± 11.7 motor units per participant were identified across post-exercise timings. In full-body condition, 34.7 ± 13.3 motor units per participant were identified in pre-exercise and an average of 18.1 ± 8.3 motor units per participant were identified across post-exercise timings.

### Effect of exercise condition on recruitment threshold and peak discharge rate

For recruitment threshold, no significant effect of exercise was observed (time effect p = 0.354, condition effect p = 0.067 and interaction effect p = 0.546) (Fig. 4A).

For peak discharge rate, there was a time effect (p = 0.002) and a condition effect (p < 0.001). Compared to pre-exercise, peak discharge rate was significantly increased from 30 seconds to 45 minutes post-exercise (all p < 0.013, d = 0.17 - 0.48). Peak discharge rate was higher in full-body condition (d = 0.09) (Fig. 4B and 4C).

**Figure 4.**
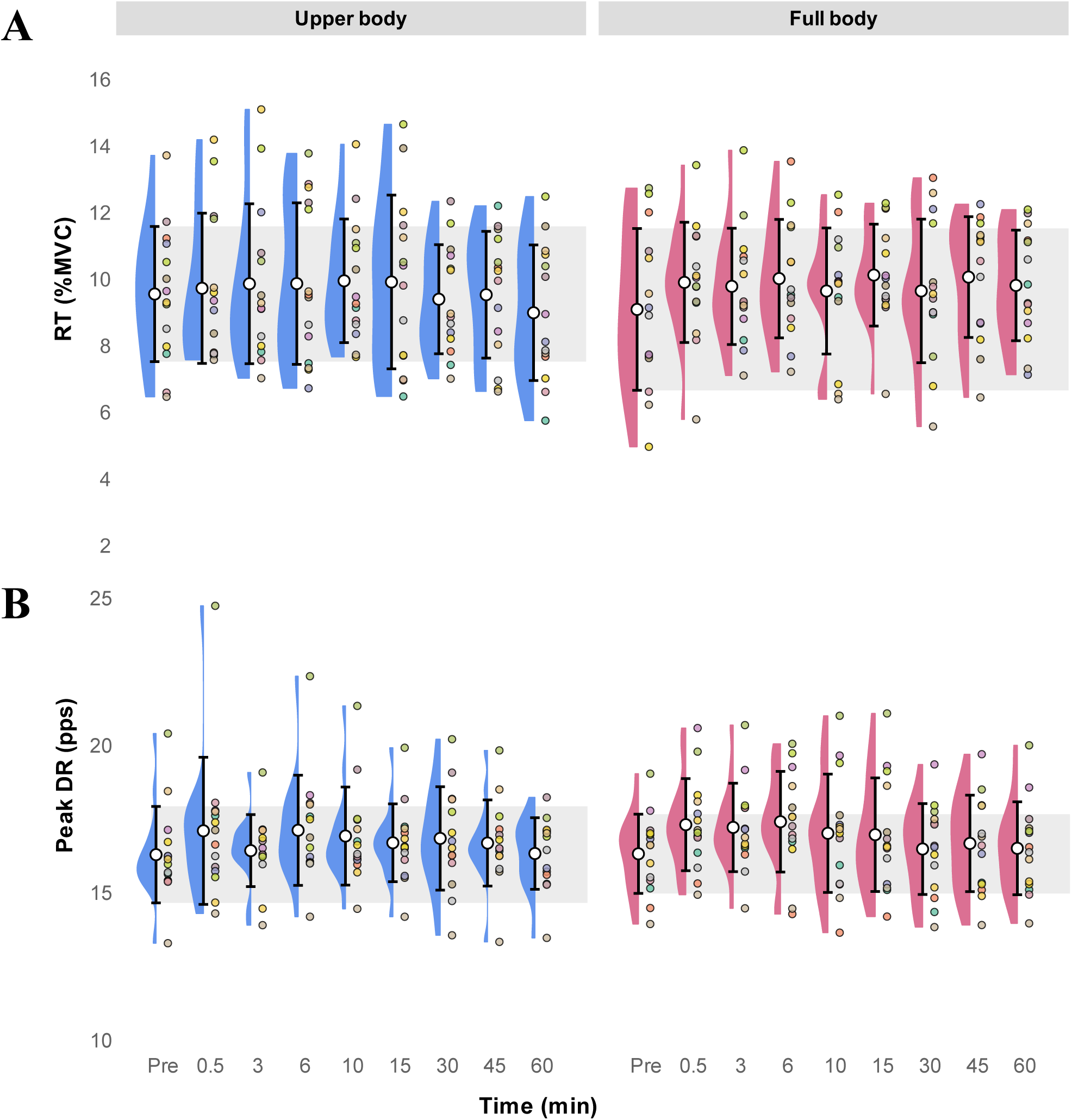
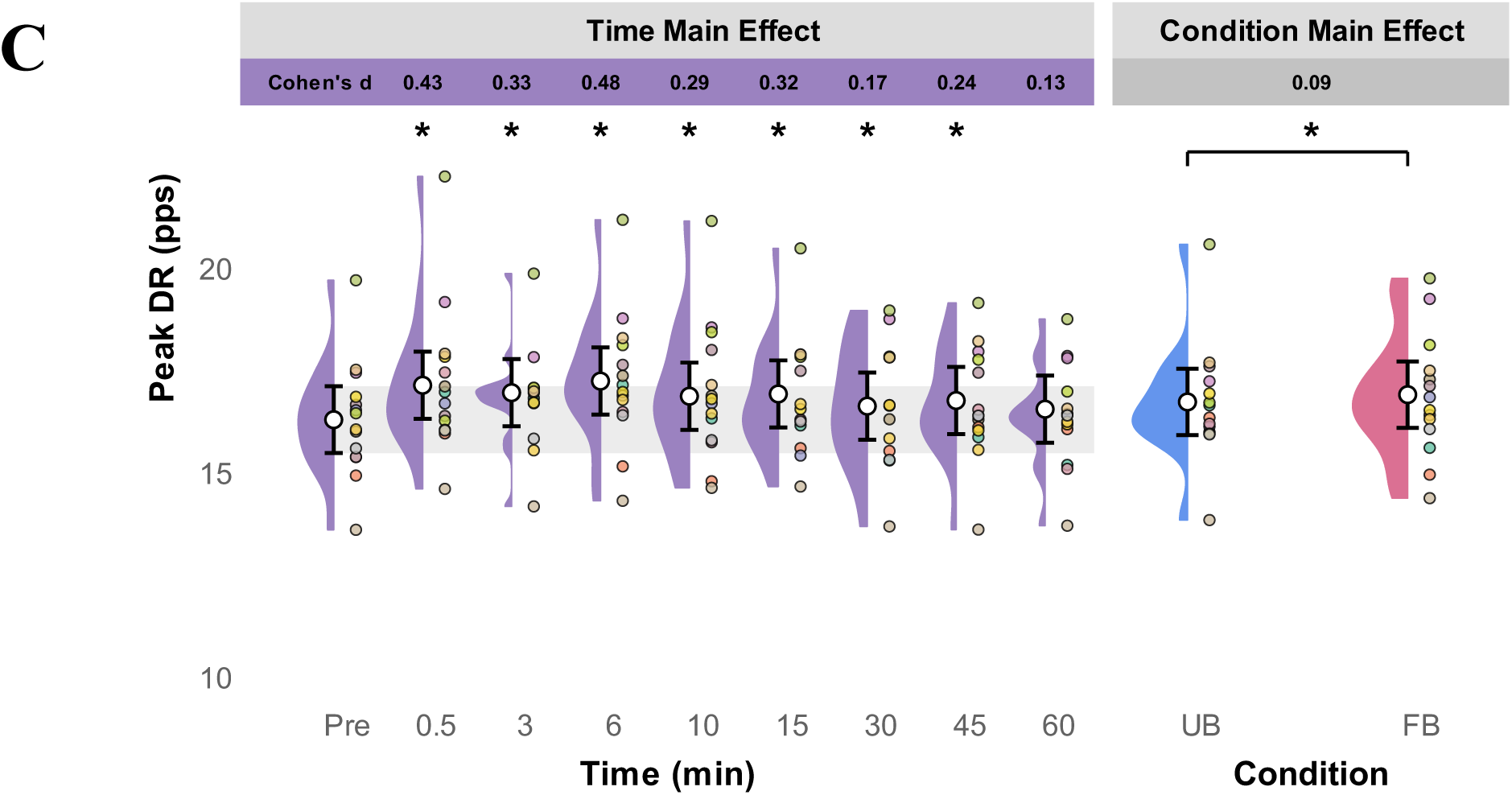
Recruitment threshold (RT) and peak discharge rates (peak DR) of motor units across timings and conditions. Recruitment threshold (*A*) and peak discharge rate (*B*) were calculated at specified timings in both conditions (upper body, in blue, full body, in pink, and averaged data, in purple). *A* and *B* show descriptive raw data displaying the time course for both conditions, while (*C*) displays linear mixed-effects model predictions for the main effects of time and condition. Colored dots represent participant averages, half-violins show the distribution of participant means, and open circles with bars represent group means (standard deviations for *A* and *B*; 95% confidence intervals for *C*). The horizontal gray shaded band represents the variability (standard deviation for *A* and *B*; 95% confidence interval for *C*) of the pre-exercise baseline. Asterisks indicate a significant difference compared to pre-exercise timing or between conditions (p < 0.05). Effect sizes (Cohen’s d) are also represented for each statistical comparison.

### Effect of exercise condition on brace height, acceleration and attenuation

For brace height, there was an interaction effect between time and condition (p < 0.001). In upper-body condition, brace height was significantly increased at 6 minutes (p < 0.001, d = 0.52), 10 minutes (p = 0.003, d = 0.38), 15 minutes (p = 0.037, d = 0.29), 45 minutes (p = 0.027, d = 0.29) and 60 minutes (p = 0.038, d = 0.28) post-exercise compared to pre-exercise. In full-body condition, brace height was significantly increased at 30 minutes (p = 0.002, d = 0.37) post-exercise compared to pre-exercise (Fig. 5A).

For acceleration, there was a time effect (p < 0.001) and a condition effect (p = 0.001). Acceleration was significantly increased at 30 seconds (p = 0.001, d = 0.29), 6 minutes (p = 0.002, d = 0.28), 10 minutes (p = 0.044, d = 0.21), 15 minutes (p < 0.001, d = 0.35), 45 minutes (p < 0.006, d = 0.23) and 60 minutes (p < 0.023, d = 0.21) post-exercise compared to pre-exercise. Acceleration was superior in full-body condition (d = 0.14) (Fig. 5B and 5C).

For attenuation, there was an interaction effect between time and condition (p < 0.001). In upper-body condition, there was no significant difference between pre- and post-exercise. In full-body condition, attenuation was significantly increased at 3 minutes (p = 0.002, d = 0.39), 6 minutes (p = 0.029, d = 0.31), 10 minutes (p = 0.050, d = 0.29) and 15 minutes (p = 0.004, d = 0.35) post-exercise (Fig. 5D).

**Figure 5.**
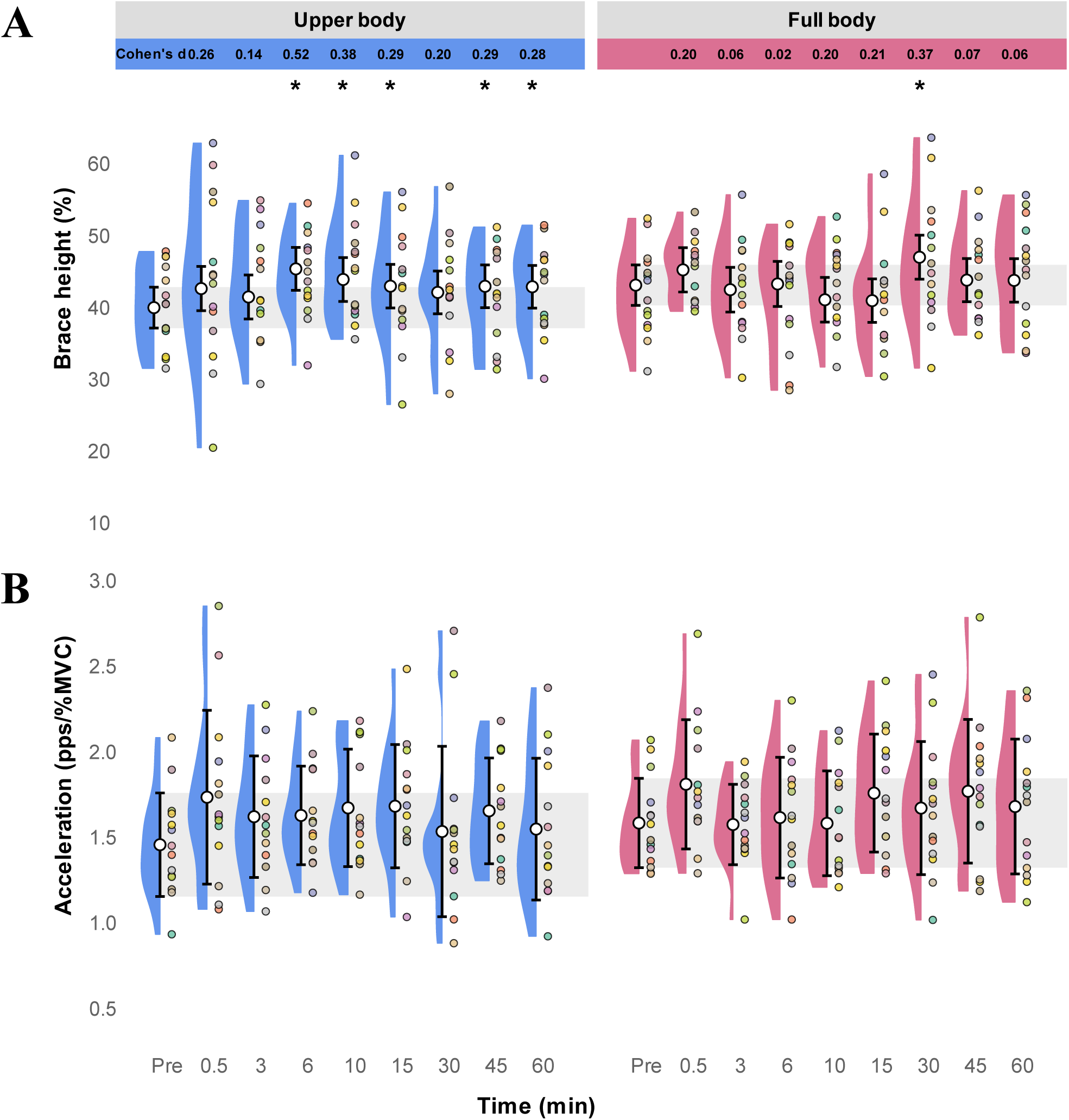
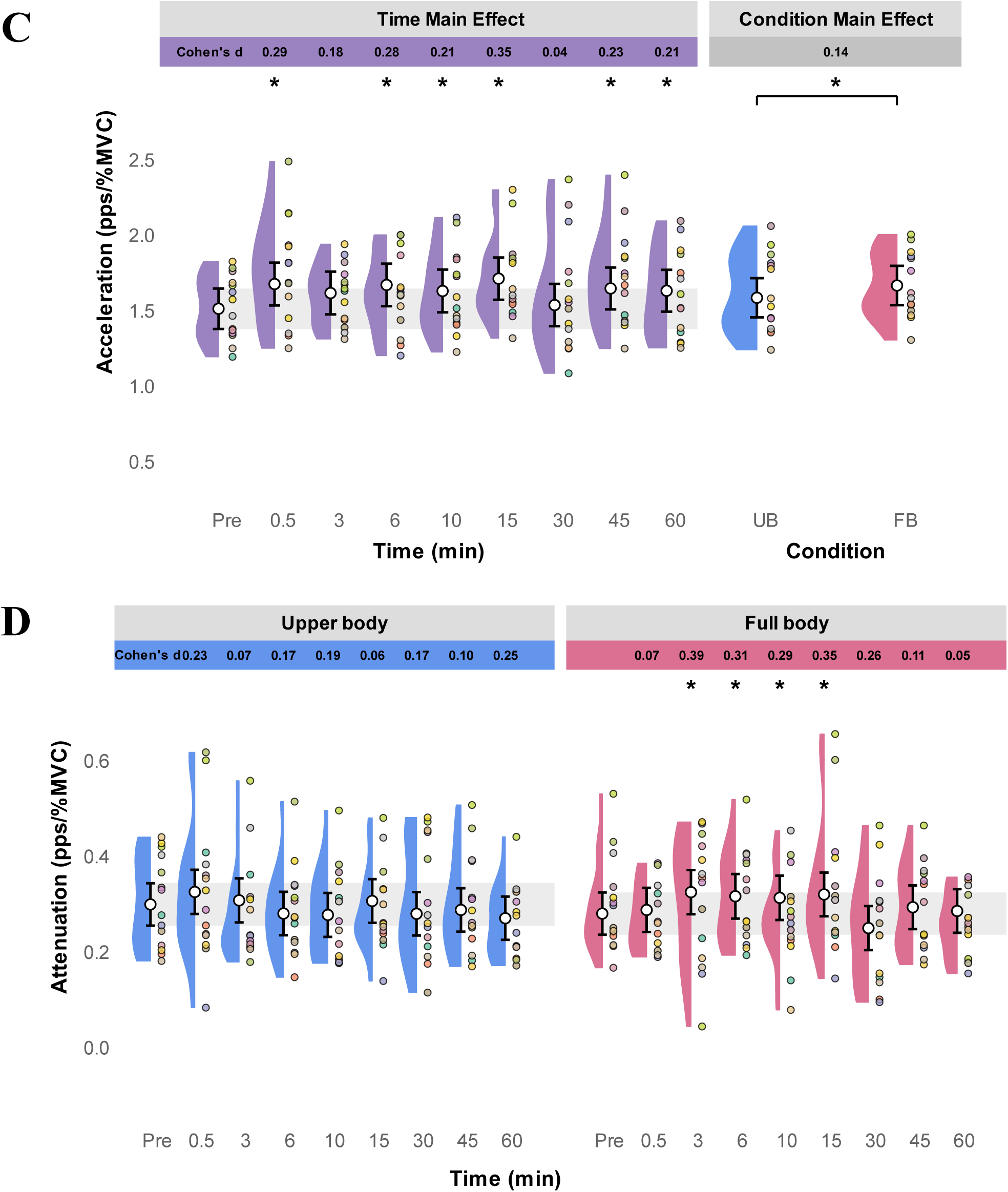
Parameters from the geometric approach of motor units across timings and conditions. Brace height (*A*), acceleration (*B, C*), and attenuation (*D*) were calculated at specified timings in both conditions (upper body, in blue, full body, in pink, and averaged data, in purple). Panels *A*, *C*, and *D* display linear mixed-effects model predictions, while panel *B* shows descriptive raw data. Colored dots represent participant averages, half-violins show the distribution of participant means, and open circles with bars represent group means (standard deviations for *B*; 95% confidence intervals for *A*, *C*, and *D*). The horizontal gray shaded band represents the variability (standard deviation for *B*; 95% confidence interval for *A*, *C* and *D*) of the pre-exercise baseline. Asterisks indicate a significant difference compared to pre-exercise timing or between conditions (p < 0.05). Effect sizes (Cohen’s d) are also represented for each statistical comparison.

### Effect of exercise condition on ΔF

For ΔF, there was a time effect (p < 0.001) and a condition effect (p < 0.001). Compared to pre-exercise, ΔF was significantly increased from 30 seconds to 60 minutes post-exercise (all p < 0.029, d = 0.20 - 0.34). ΔF was higher in upper-body condition (d = 0.21) (Fig. 6A and 6B).

**Figure 6.**
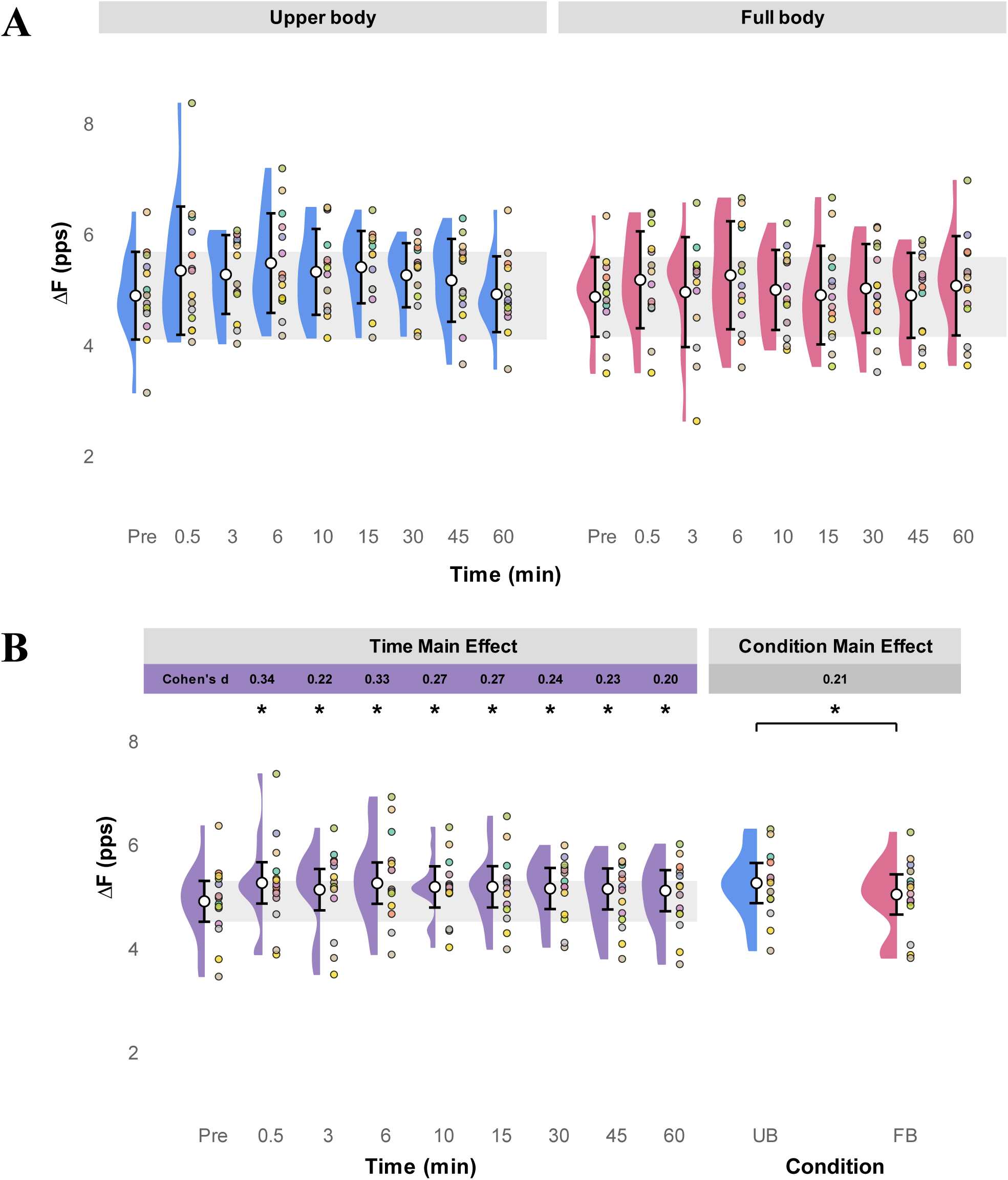
Persistent inward current estimates (ΔF) for motor units across timings and conditions. Persistent inward currents (ΔF) were calculated at specified timings in both conditions (upper body, in blue, full body, in pink, and averaged data, in purple). *A* shows descriptive raw data displaying the time course for both conditions, while *B* displays linear mixed-effects model predictions for the main effects of time and condition. Colored dots represent participant averages, half-violins show the distribution of participant means, and open circles with bars represent group means (standard deviations *A*; 95% confidence intervals *B*). The horizontal gray shaded band represents the variability (standard deviation for *A*; 95% confidence interval for *B*) of the pre-exercise baseline. Asterisks indicate a significant difference compared to pre-exercise timing or between conditions (p < 0.05). Effect sizes (Cohen’s d) are also represented for each statistical comparison.

## DISCUSSION

This study investigated how non-fatiguing physical exercise modulates human motor unit discharge pattern and the relative contributions of neuromodulatory input and excitation-inhibition balance to motoneuron output. Our findings revealed that non-fatiguing exercise led to an increase in motor unit peak discharge rate in upper-body and full-body conditions, with an effect lasting up to 45 minutes post-exercise. However, the underlying mechanisms appeared to differ depending on muscle involvement in the exercise task. When the tibialis anterior was minimally involved (i.e., upper-body condition), increases in ΔF, acceleration, and brace height suggested enhanced intrinsic motoneuron excitability through neuromodulatory input. In contrast, when the muscle was moderately involved in the task (i.e., full-body condition), the increased attenuation slope suggested a reduced contribution of inhibition with respect to excitation. Together, these findings suggest that non-fatiguing exercise induces sustained facilitation of motoneuron output through distinct spinal mechanisms, whose temporal dynamics may depend on muscle involvement.

### Both exercise condition increased motor unit peak discharge rate in the tibialis anterior

Despite differences in exercise modalities, both conditions increased motor unit peak discharge rate for up to 45 minutes post-exercise. Given that torque was matched across time points and that MVC torque remained unchanged, this higher peak discharge rate may have been offset by the recruitment of fewer motor unit to produce the same force. However, HD-EMG decomposition only identifies a subset of the active motor unit pool (Avrillon *et al*. 2024), which prevents us from confirming this hypothesis. This suggests that a single session of non-fatiguing exercise can sustainably increase motoneuron output during similar torque-matched contractions, regardless of the muscle’s involvement in the previous exercise. Consequently, these changes are unlikely to be driven by local peripheral adaptations or fatigue, pointing instead toward a central modulation of neural excitability. This aligns with reported increases in intracortical and corticospinal excitability following moderate-intensity exercise (Singh *et al*., 2014; Yamazaki et al., 2019; Youssef et al., 2024; Harroum *et al*., 2025), as well as increased motor unit peak discharge rates following remote task (Orssatto *et al*., 2022; Ugliara *et al*., 2026). Such sustained facilitation could represent a neural component of the effects of active warm-up. Importantly, increased motor unit peak discharge rate across conditions did not imply identical underlying mechanisms, as motor unit firing-pattern metrics were differentially modulated according to muscle involvement.

### Minimal muscle involvement during exercise enhances the neuromodulatory contribution to motoneuron activity

Following upper-body condition, we observed an increased contribution of PICs to the discharge pattern of tibialis anterior motor units. Specifically, the prolongation (ΔF) and the amplification (acceleration) effects of PICs were globally increased for more than 60 minutes post-exercise. Likewise, a proxy of neuromodulatory input (brace height) was increased between 6- and 60-minutes post-exercise, whereas the excitation-inhibition balance (attenuation slope) remained unchanged. Taken together, these results suggest that a 30-minute session of continuous, moderate-intensity exercise enhances the contribution of neuromodulatory input to motoneurons discharge in a muscle minimally involved in the task. This finding aligns with Nazaroff et al. (2026), who reported an increased estimated neuromodulatory input to the tibialis anterior during an acute arm-cycling task. Given the diffuse distribution of descending monoaminergic projections in the spinal cord (Heckman *et al*., 2008), such an effect likely reflects systemic monoaminergic facilitation. In particular, the serotonergic system is a plausible contributor as it is known to increase its activity in parallel with motor output (Jacobs *et al*., 2002), whereas the noradrenaline system is a less likely contributor as it is more influenced by the state of arousal (Aston-Jones *et al*., 2001). This interpretation is consistent with the hypothesis proposed by Orssatto *et al*. (2022), suggesting that acute exercise enhances the serotonin availability into the spinal cord, thereby facilitating PICs activation. Evidences from animal model align with our observations: 20-to-60 minutes of treadmill running increases the activity of serotonergic neurons in the dorsal raphe nucleus (Otsuka et *al.*, 2016; Morikawa *et al*., 2021) and the serotonin release in brain (Meeusen *et al*., 1996; Chaouloff, 1997; Gómez-Merino *et al*., 2001), with effects persisting 20-30 minutes post-exercise. Effects highlighted by these studies are all the more interesting given that central serotonergic nervous activity is correlated with cerebrospinal fluid levels of serotonin in animal (Matsumoto *et al*., 1991). Our results suggest that non-fatiguing exercise enhances neuromodulatory input in minimally involved muscles, which increases the contribution of PICs to motor unit discharge pattern. This modulation appears to persist for several dozen minutes post-exercise and likely reflects a systemic facilitation of input-output gain at the spinal motoneuron level, potentially mediated by serotonergic pathways.

### Active muscle involvement during exercise shifts the excitation-inhibition balance of **motoneurons**

In contrast to the upper-body condition, full-body rowing did not induce clear evidence for an increased neuromodulatory contribution to tibialis anterior motor unit discharge rate. Brace height, our proxy of neuromodulatory input, remained largely stable across the post-exercise period, despite a significant increase at 30 minutes post exercise. Furthermore, as ΔF and acceleration can also be influence by changes in excitation-inhibition balance (Beauchamp *et al*., 2023), their sustained increase up to 60 minutes post-exercise is difficult to interpret as a purely neuromodulatory effect. Rather, these changes may indicate a modification of the excitation-inhibition balance shaping motor unit discharge rate. Therefore, when the tibialis anterior was moderately involved in the exercise task, post-exercise modulation of motoneuron firing appeared less attributable to enhanced neuromodulatory input than in the upper-body condition.

The serotonin spillover effect could be at the origin of the absence of clear changes in neuromodulatory input after the full-body exercise. Indeed, during high levels of motor activity, the increased firing of serotonergic neurons (Jacobs *et al*., 2002) may lead to an accumulation of serotonin in the extracellular space of the spinal cord. This extrasynaptic spillover can activate inhibitory 5-HT1A receptors located on the axon initial segment of motoneurons which inhibits Na^+^ channels, responsible for the genesis of action potentials (Cotel *et al*., 2013). This inhibitory process has been proposed to act as a protective mechanism that stabilizes the motor system by counteracting excessive excitatory or neuromodulatory input during intense activity (Perrier *&* Cotel, 2015). Such an effect could therefore explain why our metric characterizing neuromodulatory input showed limited modulation in the tibialis anterior, despite a comparable cardiovascular and likely systemic load to the upper-body condition.

On the other hand, the attenuation slope increased from 3-to 15-minutes post-exercise, seeming to indicate a shift toward a reinforced push-pull organization between inhibitory and excitatory inputs, as previously described in both animal and computational models (Johnson *et al*., 2012; Johnson *et al*., 2017). In this configuration, inhibition and excitation act in opposition but in a coordinated manner, and the present increase in attenuation suggests that inhibitory input was even more reduced relative to excitation input after exercise. This potential phenomenon could therefore be the most likely cause of changes in the motoneuron firing rate after the full-body rowing exercise session. This shift in excitation-inhibition balance could also explain the similar fluctuations of ΔF and acceleration between conditions, despite the absence of brace height increase after the full-body condition. Indeed, as the contribution of PICs to motor unit discharge patterns depends on excitation-inhibition balance (Revill *&* Fuglevand, 2017; Pearcey *et al*., 2022; Goreau *et al*., 2025), a reduction in inhibitory relative to excitatory input could lead to an enhanced PICs contribution.

Taken together, these findings suggest that when a muscle is moderately involved in non-fatiguing exercise, at least two local regulatory mechanisms appear to regulate motoneuron responsiveness: First, a potentially reinforced push-pull inhibition pattern which could increase net synaptic input. Second, an inhibitory serotonergic effect which could limit PICs expression and neuromodulatory facilitation. This likely represents an adaptive strategy that increase input-output gain while preventing motoneuron hyperactivity and reducing detrimental muscle activity during and after motor activity.

### Methodological considerations

Some methodological aspects should be considered when interpreting the present findings. First, the population of motor units decomposed using HDsEMG does not represent the entire motor pool (Farina *&* Holobar, 2016). However, we were able to study a large number of motor units discharge pattern in both conditions (>1000 motor units discharge pattern in each condition), which gives us confidence in the robustness of our results. Furthermore, we did not observe any significant modifications in the recruitment thresholds in our data. As we know that variations in the recruitment threshold could also affect the reliability of quasi-geometrical approach metrics (Beauchamp *et al*., 2023), this stability allows us to confidently compare the evolution of our metrics estimating synaptic drive to motoneurons across timings in both conditions.

Although tracking the same motor units across time points would have represented the most rigorous approach to assess changes in discharge properties, this strategy was not used as the primary analysis because it substantially reduced the number of motor units available for statistical analysis. Instead, we relied on the stability of recruitment thresholds over time, which suggests that comparable motor unit populations were sampled at each time point. To address the potential risk of biased sampling of motor units, we performed a complementary analysis restricted to tracked motor units (see supplemental materials). Although this analysis was limited to effect sizes because of the reduced sample size, it confirmed the main changes identified in the full dataset, strengthening confidence in our results.

### Future directions

To clearly dissociate post-exercise systemic neuromodulation from local ionotropic or excitation-inhibition dynamics, future studies should couple motor unit discharge rate analysis with direct assessments of spinal inhibitory circuits, such as reciprocal and recurrent inhibition. Alongside these mechanistic measures, systematically manipulating exercise intensity, duration, and local muscle recruitment will be necessary to identify the exact threshold at which systemic neuromodulatory facilitation is counterbalanced by local inhibitory regulations. Furthermore, beyond the parameters of the exercise itself, it remains to be determined whether this remote facilitation is a generalized property of the motor system, or if it is restricted to muscles exhibiting intrinsically greater baseline PICs amplitudes, such as the tibialis anterior (Goreau *et al*., 2024). Ultimately, resolving these physiological boundaries in healthy cohorts will provide the foundation for extending this framework to older adults and clinical populations. Because spinal inhibition and/or monoaminergic drive are altered in some of these populations (Baudry *et al*., 2010; Crone *et al*., 2003; Hassan *et al*., 2021; Goreau *et al*., 2026), their capacity to transiently amplify motoneuronal output via non-fatiguing exercise may be modified.

## CONCLUSION

This study highlights that acute, non-fatiguing exercise induces a sustained facilitation of motoneuron output, even in a muscle minimally involved in the task. Despite this comparable facilitation, the underlying spinal adjustments differed depending on muscle involvement, reflecting either enhanced neuromodulatory input or changes in excitation-inhibition balance. Importantly, these adaptations showed different temporal patterns during recovery, suggesting that sustained increases in motoneuron output may result from distinct spinal mechanisms. These findings provide new insights into how non-fatiguing exercise modulates spinal motor control during the hour after exercise cessation, with potential implications for understanding the neuromuscular benefits of active warm-up and optimizing exercise-based rehabilitation strategies.

## DATA AVAILABILITY

The data that support the findings of this study are available on request from the corresponding author.

## SUPPLEMENTAL MATERIAL

Supplemental Material: https://doi.org/10.6084/m9.figshare.34983625.

## REFERENCES

Amann, M., Sidhu, S. K., Weavil, J. C., Mangum, T. S., & Venturelli, M. (2015). Autonomic responses to exercise: Group III/IV muscle afferents and fatigue. Autonomic Neuroscience: Basic & Clinical, 188, 19–23. 10.1016/j.autneu.2014.10.018

Aston-Jones, G., Chen, S., Zhu, Y., & Oshinsky, M. L. (2001). A neural circuit for circadian regulation of arousal. Nature Neuroscience, 4(7), 732–738.

Astrand, P. O., & Ryhming, I. (1954). A nomogram for calculation of aerobic capacity (physical fitness) from pulse rate during sub-maximal work. Journal of Applied Physiology, 7(2), 218–221. 10.1152/jappl.1954.7.2.218

Avrillon, S., Hug, F., Baker, S. N., Gibbs, C., & Farina, D. (2024). Tutorial on MUedit: An open-source software for identifying and analysing the discharge timing of motor units from electromyographic signals. Journal of Electromyography and Kinesiology, 77, 102886. 10.1016/j.jelekin.2024.102886

Baudry, S., Maerz, A. H., & Enoka, R. M. (2010). Presynaptic modulation of Ia afferents in young and old adults when performing force and position control. Journal of Neurophysiology, 103(2), 623–631. 10.1152/jn.00839.2009

Beauchamp, J. A., Khurram, O. U., Dewald, J. P. A., Heckman, C., & Pearcey, G. E. P. (2022). A computational approach for generating continuous estimates of motor unit discharge rates and visualizing population discharge characteristics. Journal of Neural Engineering, 19(1), 016007. 10.1088/1741-2552/ac4594

Beauchamp, J. A., Pearcey, G. E. P., Khurram, O. U., Chardon, M., Wang, Y. C., Powers, R. K., Dewald, J. P. A., & Heckman, C. (2023). A geometric approach to quantifying the neuromodulatory effects of persistent inward currents on individual motor unit discharge patterns. Journal of Neural Engineering, 20(1), 016034. 10.1088/1741-2552/acb1d7

Chardon, M. K., Wang, Y. C., Garcia, M., Besler, E., Beauchamp, J. A., D’Mello, M., Powers, R. K., & Heckman, C. J. (2024). Supercomputer framework for reverse engineering firing patterns of neuron populations to identify their synaptic inputs. eLife, 12, Article RP90624. 10.7554/eLife.90624.3

Cotel, F., Exley, R., Cragg, S. J., & Perrier, J. F. (2013). Serotonin spillover onto the axon initial segment of motoneurons induces central fatigue by inhibiting action potential initiation. Proceedings of the National Academy of Sciences, 110(12), 4774–4779. 10.1073/pnas.1216150110

Crone, C., Johnsen, L. L., Biering-Sørensen, F., & Nielsen, J. B. (2003). Appearance of reciprocal facilitation of ankle extensors from ankle flexors in patients with stroke or spinal cord injury. Brain, 126(2), 495–507. 10.1093/brain/awg036

Del Vecchio, A., Holobar, A., Falla, D., Felici, F., Enoka, R. M., & Farina, D. (2020). Tutorial: Analysis of motor unit discharge characteristics from high-density surface EMG signals. Journal of Electromyography and Kinesiology, 53, 102426. 10.1016/j.jelekin.2020.102426

Farina, D., & Holobar, A. (2016). Characterization of human motor units from surface EMG decomposition. Proceedings of the IEEE, 104(2), 353–373.

Gomez-Merino, D., Béquet, F., Berthelot, M., Chennaoui, M., & Guezennec, C. Y. (2001). Site-dependent effects of an acute intensive exercise on extracellular 5-HT and 5-HIAA levels in rat brain. Neuroscience letters, 301(2), 143–146.

Gorassini, M., Yang, J. F., Siu, M., & Bennett, D. J. (2002). Intrinsic activation of human motoneurons: Possible contribution to motor unit excitation. Journal of Neurophysiology, 87(4), 1850–1858. 10.1152/jn.00024.2001

Goreau, V., Hug, F., Jannou, A., Dernoncourt, F., Crouzier, M., & Cattagni, T. (2024). Estimates of persistent inward currents in lower limb muscles are not different between inactive, resistance-trained, and endurance-trained young males. Journal of Neurophysiology, 131(2), 166–175. 10.1152/jn.00278.2023

Goreau, V., Morvan, Q., Hug, F., Le Sant, G., Gross, R., & Cattagni, T. (2025). Modulation of persistent inward currents in alpha motoneurons with joint angle depends on muscle length. Journal of Neurophysiology. Advance online publication. 10.1152/jn.00097.2025

Goreau, V., Hug, F., Simon, L., Le Sant, G., Gross, R., & Cattagni, T. (2026). Disrupted inhibitory-excitatory balance underlies spinal motoneuron dysfunction in incomplete spinal cord injury. bioRxiv. 10.64898/2026.02.14.705880

Harroum, N., O’Farrell, A., Youssef, L., Bohbot, L., Maati, H., Joubert, M., Pageaux, B., & Neva, J. L. (2025). Motor cortical excitability and inhibition are differentially modulated by aerobic exercise intensity. Cerebral Cortex, 35(8), Article bhaf230. 10.1093/cercor/bhaf230

Hassan, A., Thompson, C. K., Negro, F., Cummings, M., Powers, R. K., Heckman, C. J., Dewald, J. P. A., & McPherson, L. M. (2020). Impact of parameter selection on estimates of motoneuron excitability using paired motor unit analysis. Journal of Neural Engineering, 17(1), 016063. 10.1088/1741-2552/ab5eda

Hassan, A. S., Fajardo, M. E., Cummings, M., McPherson, L. M., Negro, F., Dewald, J. P. A., Heckman, C. J., & Pearcey, G. E. P. (2021). Estimates of persistent inward currents are reduced in upper limb motor units of older adults. The Journal of Physiology, 599(21), 4865–4882. 10.1113/JP282063

Heckman, C. J., Gorassini, M. A., & Bennett, D. J. (2005). Persistent inward currents in motoneuron dendrites: Implications for motor output. Muscle & Nerve, 31(2), 135–156. 10.1002/mus.20261

Heckman, C. J., Johnson, M., Mottram, C., & Schuster, J. (2008). Persistent inward currents in spinal motoneurons and their influence on human motoneuron firing patterns. The Neuroscientist, 14(3), 264–275. 10.1177/1073858408314986

Hug, F., Avrillon, S., Del Vecchio, A., Casolo, A., Ibanez, J., Nuccio, S., Rossato, J., Holobar, A., & Farina, D. (2021). Analysis of motor unit spike trains estimated from high-density surface electromyography is highly reliable across operators. Journal of Electromyography and Kinesiology, 58, 102548. 10.1016/j.jelekin.2021.102548

Jacobs, B. L., Martín-Cora, F. J., & Fornal, C. A. (2002). Activity of medullary serotonergic neurons in freely moving animals. Brain Research Reviews, 40(1-3), 45–52.

Johnson, M. D., Hyngstrom, A. S., Manuel, M., & Heckman, C. J. (2012). Push-pull control of motor output. The Journal of Neuroscience, 32(13), 4592–4599. 10.1523/JNEUROSCI.4709-11.2012

Johnson, M. D., Thompson, C. K., Tysseling, V. M., Powers, R. K., & Heckman, C. J. (2017). The potential for understanding the synaptic organization of human motor commands via the firing patterns of motoneurons. Journal of Neurophysiology, 118(1), 520–531.

Kuznetsova, A., Brockhoff, P. B., & Christensen, R. H. B. (2017). lmerTest package: Tests in linear mixed effects models. Journal of Statistical Software, 82(13). 10.18637/jss.v082.i13

Li, X., & Bennett, D. J. (2007). Apamin-sensitive calcium-activated potassium currents (SK) are activated by persistent calcium currents in rat motoneurons. Journal of Neurophysiology, 97(5), 3314–3330. 10.1152/jn.01068.2006

Mackay, K., Orssatto, L. B. R., Polman, R., Van Der Pols, J. C., & Trajano, G. S. (2023). Caffeine does not influence persistent inward current contribution to motoneuron firing. Journal of Neurophysiology, 130(6), 1529–1540. 10.1152/jn.00350.2023

Matsumoto, M., Togashi, H., Yoshioka, M., Morii, K., Hirokami, M., Tochihara, M., Ikeda, T., Saito, Y., & Saito, H. (1991). Significant correlation between cerebrospinal fluid and brain levels of norepinephrine, serotonin and acetylcholine in anesthetized rats. Life Sciences, 48(8), 823–829. 10.1016/0024-3205(91)90098-v

Meeusen, R., Thorré, K., Chaouloff, F., Sarre, S., De Meirleir, K., Ebinger, G., & Michotte, Y. (1996). Effects of tryptophan and/or acute running on extracellular 5-HT and 5-HIAA levels in the hippocampus of food-deprived rats. Brain Research, 740(1-2), 245–252. 10.1016/s0006-8993(96)00872-4

Morikawa, R., Kubota, N., Amemiya, S., Nishijima, T., & Kita, I. (2021). Interaction between intensity and duration of acute exercise on neuronal activity associated with depression-related behavior in rats. The Journal of Physiological Sciences, 71, Article 1. 10.1186/s12576-020-00788-5

Motl, R. W., & Dishman, R. K. (2003). Acute leg-cycling exercise attenuates the H-reflex recorded in soleus but not flexor carpi radialis. Muscle & Nerve, 28(5), 609–614. 10.1002/mus.10479

Motl, R. W., Knowles, B. D., & Dishman, R. K. (2003). Acute bouts of active and passive leg cycling attenuate the amplitude of the soleus H-reflex in humans. Neuroscience Letters, 347(2), 69–72. 10.1016/s0304-3940(03)00652-9

Nazaroff, B. M., Mitchell, E. R., & Pearcey, G. E. (2026). Arm locomotor behaviour affects motor unit discharge characteristics in the stationary leg. bioRxiv. 10.64898/2026.08.17.745355

Negro, F., Muceli, S., Castronovo, A. M., Holobar, A., & Farina, D. (2016). Multichannel intramuscular and surface EMG decomposition by convolutive blind source separation. Journal of Neural Engineering, 13(2), 026027. 10.1088/1741-2560/13/2/026027

Orssatto, L. B. R., Fernandes, G. L., Blazevich, A. J., & Trajano, G. S. (2022). Facilitation-inhibition control of motor neuronal persistent inward currents in young and older adults. The Journal of Physiology, 600(23), 5101–5117. 10.1113/JP283708

Otsuka, T., Nishii, A., Amemiya, S., Kubota, N., Nishijima, T., & Kita, I. (2016). Effects of acute treadmill running at different intensities on activities of serotonin and corticotropin-releasing factor neurons, and anxiety-and depressive-like behaviors in rats. Behavioural Brain Research, 298(Part B), 44–51. 10.1016/j.bbr.2015.10.055

Perrier, J. F., & Cotel, F. (2015). Serotonergic modulation of spinal motor control. Current Opinion in Neurobiology, 33, 1–7. 10.1016/j.conb.2014.12.008

Pearcey, G. E., Khurram, O. U., Beauchamp, J. A., Negro, F., & Heckman, C. J. (2022). Antagonist tendon vibration dampens estimates of persistent inward currents in motor units of the human lower limb. bioRxiv.

Pierrot-Deseilligny, E., & Burke, D. (2012). The circuitry of the human spinal cord: Spinal and corticospinal mechanisms of movement. Cambridge University Press.

Powers, R. K., & Binder, M. D. (2001). Input-output functions of mammalian motoneurons. Reviews of Physiology, Biochemistry and Pharmacology, 143, 137–263. 10.1007/BFb0115594

Powers, R. K., & Heckman, C. J. (2015). Contribution of intrinsic motoneuron properties to discharge hysteresis and its estimation based on paired motor unit recordings: A simulation study. Journal of Neurophysiology, 114(1), 184–198. 10.1152/jn.00019.2015

Revill, A. L., & Fuglevand, A. J. (2017). Inhibition linearizes firing rate responses in human motor units: Implications for the role of persistent inward currents. The Journal of Physiology, 595(1), 179–191. 10.1113/JP272823

Sherrington, C. S. (1906). The integrative action of the nervous system. Yale University Press.

Singh, A. M., Duncan, R. E., Neva, J. L., & Staines, W. R. (2014). Aerobic exercise modulates intracortical inhibition and facilitation in a nonexercised upper limb muscle. BMC Sports Science, Medicine and Rehabilitation, 6, Article 23. 10.1186/2052-1847-6-23

Stephenson, J. L., & Maluf, K. S. (2011). Dependence of the paired motor unit analysis on motor unit discharge characteristics in the human tibialis anterior muscle. Journal of Neuroscience Methods, 198(1), 84–92. 10.1016/j.jneumeth.2011.03.018

Ugliara, L., Orssatto, L. B. R., Vieira, A., & Trajano, G. S. (2026). Isometric handgrip contraction increases tibialis anterior intrinsic motoneuron excitability in a dose-dependent manner. Experimental Physiology. Advance online publication. 10.1113/EP092961

Yamazaki, Y., Sato, D., Yamashiro, K., Nakano, S., Onishi, H., & Maruyama, A. (2019). Acute low-intensity aerobic exercise modulates intracortical inhibitory and excitatory circuits in an exercised and a non-exercised muscle in the primary motor cortex. Frontiers in Physiology, 10, Article 1361. 10.3389/fphys.2019.01361

Youssef, L., Harroum, N., Francisco, B. A., Johnson, L., Arvisais, D., Pageaux, B., … Neva, J. L. (2024). Neurophysiological effects of acute aerobic exercise in young adults: A systematic review and meta-analysis. Neuroscienc

